# Gα_q_ modulates the energy metabolism of osteoclasts

**DOI:** 10.1101/2022.04.06.487374

**Authors:** S. Chakraborty, B. Handrick, D. Yu, K. Bode, A. Hafner, J. Schenz, D. Schaack, F. Uhle, T. Tachibana, S. Kamitani, T. Vogl, K.F. Kubatzky

**Affiliations:** Department of Infectious Diseases, Medical Microbiology and Hygiene, Heidelberg University Hospital, Im Neuenheimer Feld 324, 69120 Heidelberg, Germany; Department of Transplant Immunology and Immunogenetics, All India Institute of Medical Sciences, New Delhi 1100029, India; Department of Anesthesiology, Heidelberg University Hospital, Im Neuenheimer Feld 420, 69120 Heidelberg, Germany; Department of Chemistry and Bioengineering, Graduate School of Engineering, Osaka Metropolitan University, Japan; Department of Nutrition, Graduate School of Human Life and Ecology, Osaka Metropolitan University, Japan; Institute of Immunology, University Hospital Münster, Röntgenstr.21, 48149 Münster, Germany

**Author notes:** **corresponding author:** E-mail address (K.F. Kubatzky), @kkubatzky. Dept. Molecular Diagnostics, Laboratory Dr. Limbach and Colleagues, Im Breitspiel 16, 69126 Heidelberg, Germany.

**Keywords:** immunometabolism, Gαq, osteoclast, *Pasteurella multocida* toxin, mitochondria, OPA1, STAT3, rheumatoid arthritis

## Abstract

The bacterial protein toxin Pasteurella multocida toxin (PMT) mediates RANKL-independent osteoclast differentiation. Although these osteoclasts are small, their resorptive activity is high and destroys the nasal turbinate bones of pigs. Analysis of the proteome of classical and toxin-derived osteoclasts showed that PMT induces the upregulation of metabolic pathways. This includes strong glycolytic activity, increased expression of GLUT1 and upregulation of the mTOR pathway. As OxPhos components are also expressed more efficiently, cells display increased mitochondrial respiration. We found that the heterotrimeric G protein Gαq plays a central role in this hypermetabolic cell activation. Gαq triggers mitochondrial relocalisation of pSerSTAT3 and an increase in OPA1 expression. Overexpression of Gαq in Hoxb8 cells mimicked this hypermetabolic phenotype and resulted in higher glycolytic and mitochondrial activity as well as increased bone resorptive activity. Rheumatoid arthritis patients show an increase in *Gnaq* expression especially in the synovial fluid, suggesting that Gαq is a target of pathophysiological relevance.

## Introduction

Bone remodelling is a continuous process throughout adult life and various soluble factors and cells influence this process. Changes in the microenvironment of bone due to infection or inflammatory processes lead to bone loss due to excessive formation and function of osteoclast (OC). OC are bone resorbing haematopoietic cells that can be differentiated from their respective monocyte precursor cells through the cytokines macrophage colony stimulating factor (M-CSF) and receptor activator of NF-kB ligand (RANKL). Pro-inflammatory cytokines such as IL-6, TNF-α and IL-1β can act as additional stimuli that enhance OC formation (Shaw and Gravallese, 2016). Therefore, auto-inflammatory pathologies such as rheumatoid arthritis or chronic inflammation, e.g. osteomyelitis, are often accompanied by a decrease in bone density. During differentiation, preosteoclasts fuse and form polycaryons before becoming mature OC. OC are defined as multi-nucleated cells that stain tartrate resistant acidic phosphatase (TRAP)-positive and resorb bone. To enable cellular fusion and bone resorption, OC increase their mitochondria to meet the energy demand (Arnett and Orriss, 2017). However, our understanding of the changes in metabolic activity during OC differentiation is still limited. In addition to TCA cycle activation and OxPhos activity, glycolysis is involved and glucose transporter GLUT1 is upregulated in a RANKL-dependent manner (Indo et al., 2013). The importance of mitochondrial activity is highlighted by the finding that inhibition of mitochondria through deletion of the complex I component Ndufs4 prevents OC differentiation and shifts macrophages towards M1 activation and inflammation (Jin et al., 2014). Due to the close relationship between macrophages and OC it can be expected that changes in macrophage activity, for example during bacterial bone infections or auto-inflammatory diseases, will impact the ability of macrophages to differentiate into OC and shape the activity of the resulting OC, respectively. The possibility to manipulate cell metabolism is by now well-established in cancer therapy, but has only recently been discovered as an interesting option in diseases like RA, systemic lupus erythematosus or osteoarthritis (Mao et al., 2020; Onuora, 2016; Sanchez-Lopez et al., 2019; Takeshima et al., 2019).

*Pasteurella multocida* toxin (PMT), a protein toxin causing atrophic rhinitis in pigs and the severe destruction of nasal bone, is produced by toxigenic *Pasteurella multocida* strains (Kubatzky, 2012). Deamidation and subsequent activation of heterotrimeric G proteins causes cytoskeletal rearrangement, proliferation, protection from apoptosis and the differentiation of macrophages into OC (Kubatzky et al., 2013). PMT causes osteoclastogenesis RANKL-independently through Gαq-dependent activation of nuclear factor of activated T cells c1 (NFATc1) and NF-kB-mediated production of pro-inflammatory cytokines (Chakraborty et al., 2017). The importance of heterotrimeric G proteins for bone formation only recently got considered. Gα13 is a negative regulator of OC fusion that restricts OC size (Nakano et al., 2019; Wu et al., 2017), whereas Gα12 enhances OC formation and activity by positively regulating NFATc1 signalling and bone resorption (Song et al., 2018). The role of Gαq in osteoclastogenesis has not been studied. However, GPR91, the receptor for succinate, which signals via Gαq, was recently shown to be important in pathological bone loss in rheumatoid arthritis (Littlewood-Evans et al., 2016). As PMT is a strong activator of Gαq, we used PMT as a tool to investigate the role of mitochondrial Gαq during OC differentiation.

Our proteome analysis of classical and toxin-derived OCs showed an overrepresentation of genes involved in glycolysis and metabolic pathways in PMT-stimulated samples which was verified by Seahorse analyses of glycolysis and extracellular acidification (ECAR) measurements. Overexpression of Gαq is central for serine phosphorylation and mitochondrial localisation the of signal transducer and activator of transcription (STAT3) and increased expression of the large dynamin-like GTPase OPA1 resulting in more efficient mitochondrial cristae formation, enhanced ETC expression and increased resorptive activity.

In summary, we show that Gαq plays an important role in the induction of a hypermetabolic phenotype in macrophages characterized by an upregulation of glycolytic activity as well as an enhanced capacity for oxygen consumption facilitating osteoclast differentiation and activity. Enhanced expression of Gαq was also observed in cells from synovial fluid of (RA) patients indicating its influence in disease pathogenesis.

## Results

### Distinct proteomes between PMT and MCSF/RANKL induced osteoclasts

We recently investigated the mechanism of PMT-induced osteoclastogenesis from bone marrow-derived macrophages. Despite the striking difference in cellular morphology, the functional ability of PMT-derived and classical OC to resorb bone is comparable (Chakraborty *et al*., 2017). Generally reduced fusion is linked to ineffective resorption (Mizoguchi et al., 2013), but the drastic effects of PMT in pigs suffering from atrophic rhinitis argue against this (Wilkie et al., 2012). We therefore tried to understand how PMT OCs compensate for the loss of fusogenic activity and performed an analysis of the OC proteome (supp Fig 1A). For an initial analysis the spots that were most highly upregulated by PMT were identified and analysed by mass spectrometry. Performing a GO enrichment analysis showed that metabolic pathways, the biosynthesis of amino acids as well as glycolysis and gluconeogenesis events (Panther analysis of KEGG pathways) were significantly overrepresented after PMT treatment (Fig 1A, supp. Fig 1B and table 1).

**Table 1.**
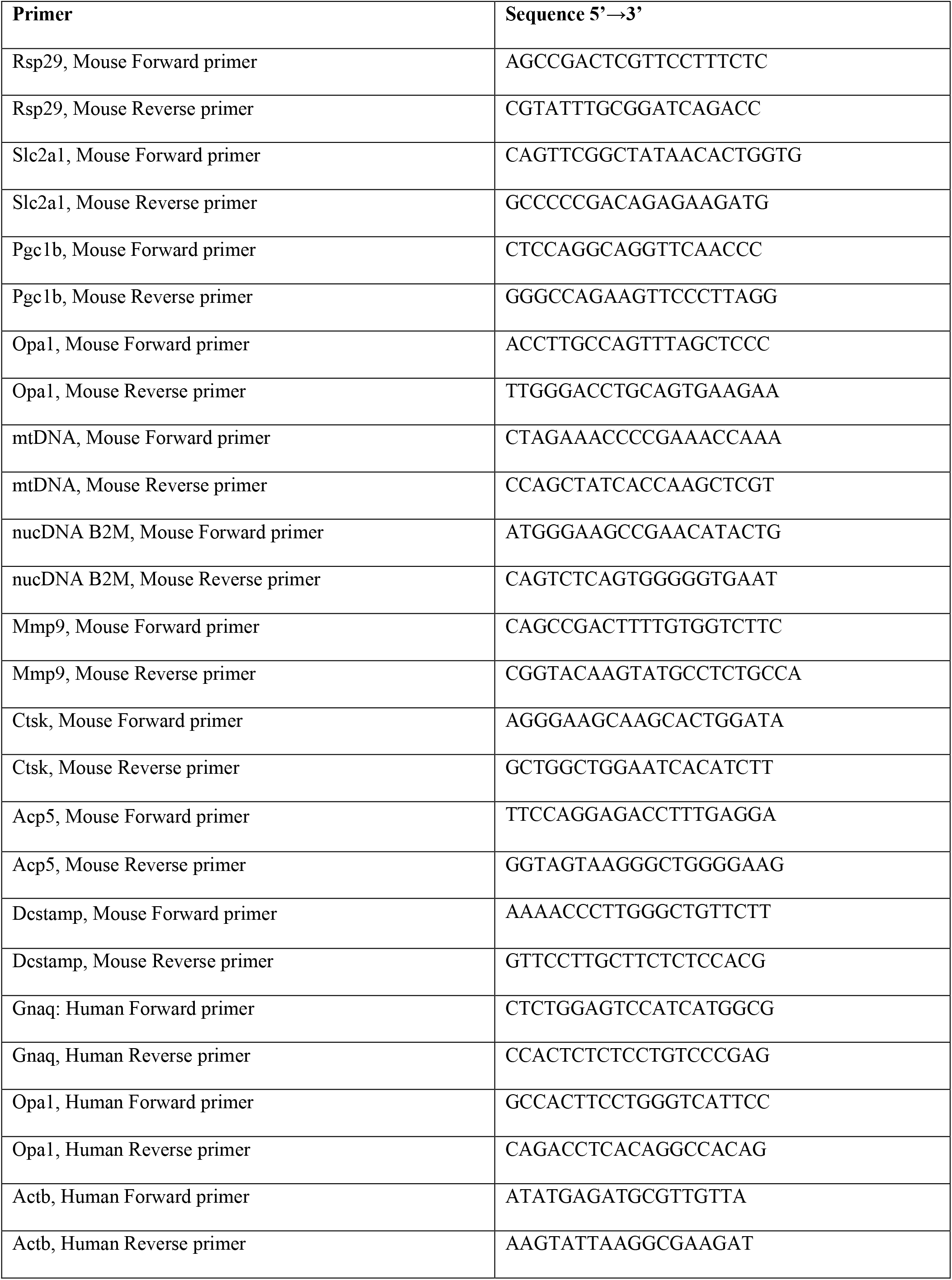
Primer;

**Figure 1:**
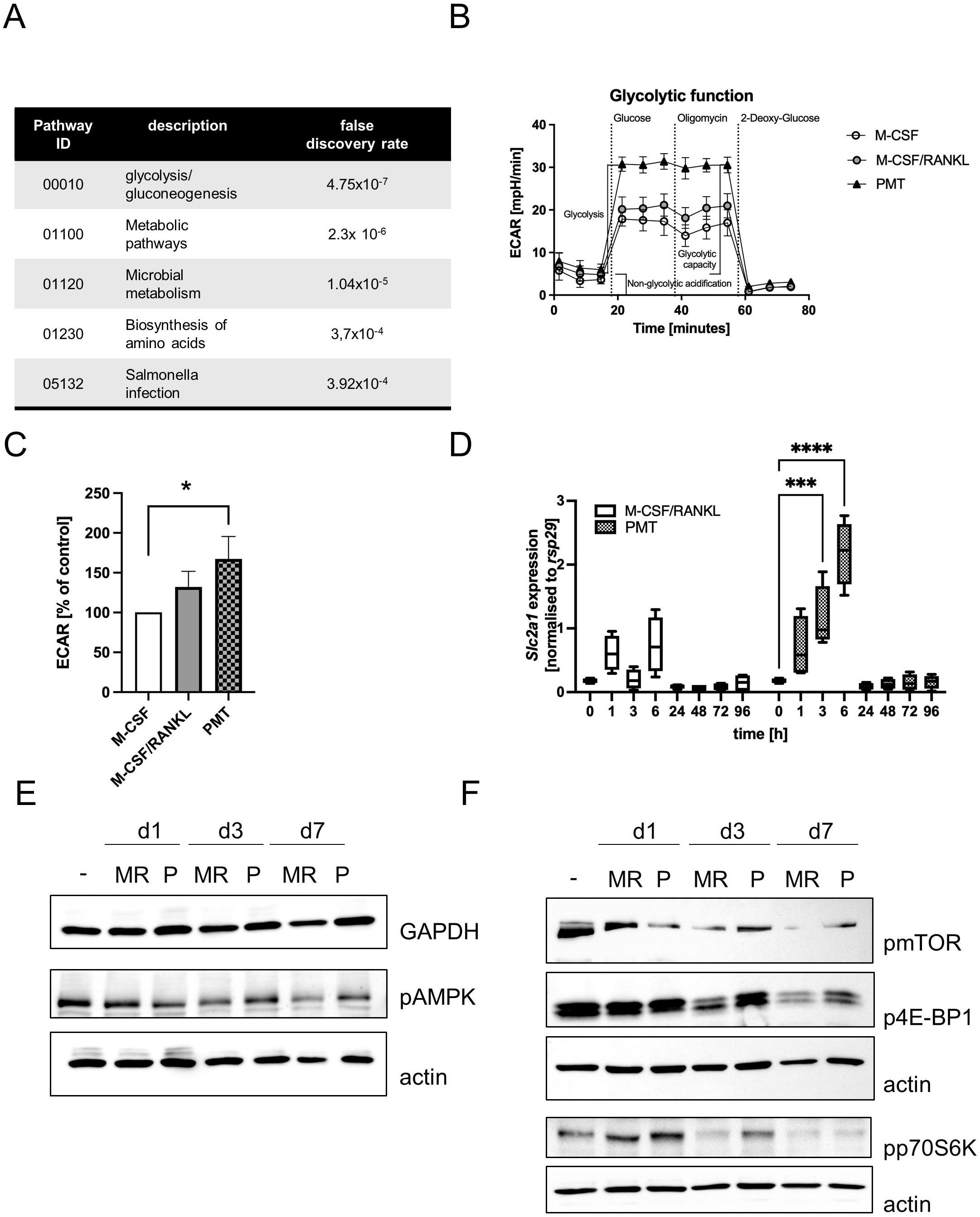
Comparison of osteoclast glycolytic activities during differentiation. (A) Proteins that were upregulated with PMT (10 most highly picked, results for 7 obtained) were subjected to a Panther GO analysis. (B)(C) Glycolysis of BMDM differentiated for 3 days with M-CSF, M-CSF/RANKL or PMT was investigated in a Seahorse glycolysis assay with quantification of ECAR (n=3). ECAR results were normalised to the M-CSF-treated control. For statistical analysis a Friedman test was used on raw data. (D) BMDM were treated with M-CSF/RANKL or PMT and samples were taken at the indicated time-points to quantify the expression of *Slc2a1* by RT-PCR (n=4). Statistical analysis was performed by 2-way ANOVA. (E)(F) BMDM were stimulated as indicated and lysates from day 0, 1, 3 and 7 were probed for the expression of pAMPK and GAPDH (E) or pmTOR, p4E-BP1, pp70S6K, respectively (F). Actin was used as a lysate control (n=4).

### Differential regulation of glycolytic pathways

To verify the computational results, we analysed the glycolytic pathway and performed a Seahorse glycolytic stress test assay (Fig 1B). Indeed, in PMT-treated cells, the extracellular acidification rate (ECAR) increased significantly after glucose addition compared to M-CSF treated cells, corresponding to higher glycolysis (Fig 1C). M-CSF/RANKL-treated OCs also show enhanced glycolytic activity, although less prominent than PMT. This was corroborated by our finding that the glucose transporter *Slc2a1* transcription was induced at early time points by PMT but only weakly by M-CSF/RANKL (Fig 1D). Because an increase in glycolysis is associated with anabolic pathways and a subsequent decrease in intracellular ATP levels, we investigated the expression of glyceraldehyde-3-phosphate dehydrogenase (GAPDH) and activation of AMPK. As suggested by the initial proteomic data, GAPDH was expressed stronger in PMT-treated cells (Fig 1E). RANKL can induce AMPK activation through phosphorylation on Thr172 of its alpha-subunit (Lee et al., 2010). However, while pAMPK decreased during classical osteoclastogenesis, pAMPK levels were higher im PMT-treated cells on days 3 and 7.

In RAW 264.7 macrophages the mTOR pathway is essential for PMT-induced OC formation, but dispensable for cytokine production and cellular viability (Kloos et al., 2015). We investigated the ability of PMT to activate the mTOR pathway, as mTOR is a central switch that regulates cellular metabolic activity (Fig 1F). Compared to the untreated control, mTOR shifted to a higher molecular weight in M-CSF/RANKL and PMT-treated samples suggesting post-translational modification. The levels of mTOR were increased in M-CSF/RANKL treated cells on day 1, while PMT maintained the high expression until day 7. Both downstream effector molecules, 4EB-P1 and p70S6K were phosphorylated in PMT-treated cells at all time-points, whereas M-CSF/RANKL-treated cells only showed high phosphorylation levels compared to untreated cells on day1.

### Mitochondrial activity during osteoclastogenesis

Mitochondrial biogenesis is an important part of RANKL-mediated OC differentiation that requires RelB and p65 (Zeng et al., 2015). Figure 2A shows that in classical and toxin-derived OC, RelB was strongly activated at an early timepoint (d 1) while PMT-treated cells also showed strong RelB expression at later time-points. Activation of the canonical NF-kB pathway was found in both OC types, but again the increase in p65 phosphorylation was stronger in PMT-treated cells. Consequently, Peroxisome proliferator-activated receptor gamma coactivator 1-beta (*Pgc1b*) transcription, the gene encoding a transcription factor responsible for mitochondrial biogenesis, was upregulated in both OC, although statistical significance was not reached (Fig 2B). Therefore, we observed an increase in the mitochondrial copy number in PMT-treated and M-CSF/RANKL treated cells with no significant difference between PMT- and M-CSF/RANKL-treated cells (Fig 2C). We measured the activity of mitochondria with Mitotracker Deep Red FM and found that PMT OC showed a stronger mitochondrial activity at all time points (Fig 2D). To understand this, we analysed the expression of OxPhos proteins and found a prominent upregulation of ETC complexes especially by PMT treatment (Fig 2E). Morita et al. had reported that the mTOR-activated translation initiation factor elF4 is a central mediator of the translation of ETC components (Morita et al., 2015). This prevents that mTOR-mediated glycolysis and anabolic activity will eventually deprive the cell of ATP and allows the cell to switch back again to OxPhos when needed. This is also reflected by the result shown in Figure 1F. The higher phosphorylation and subsequent inactivation of p4-EBP1 by PMT leads to the initiation of elF4-mediated translation and a more pronounced increase in protein translation. As seen in a metabolic pulse chase experiment with the methionine analogue HPG coupled to Alexa488 (Fig 2F).

**Figure 2:**
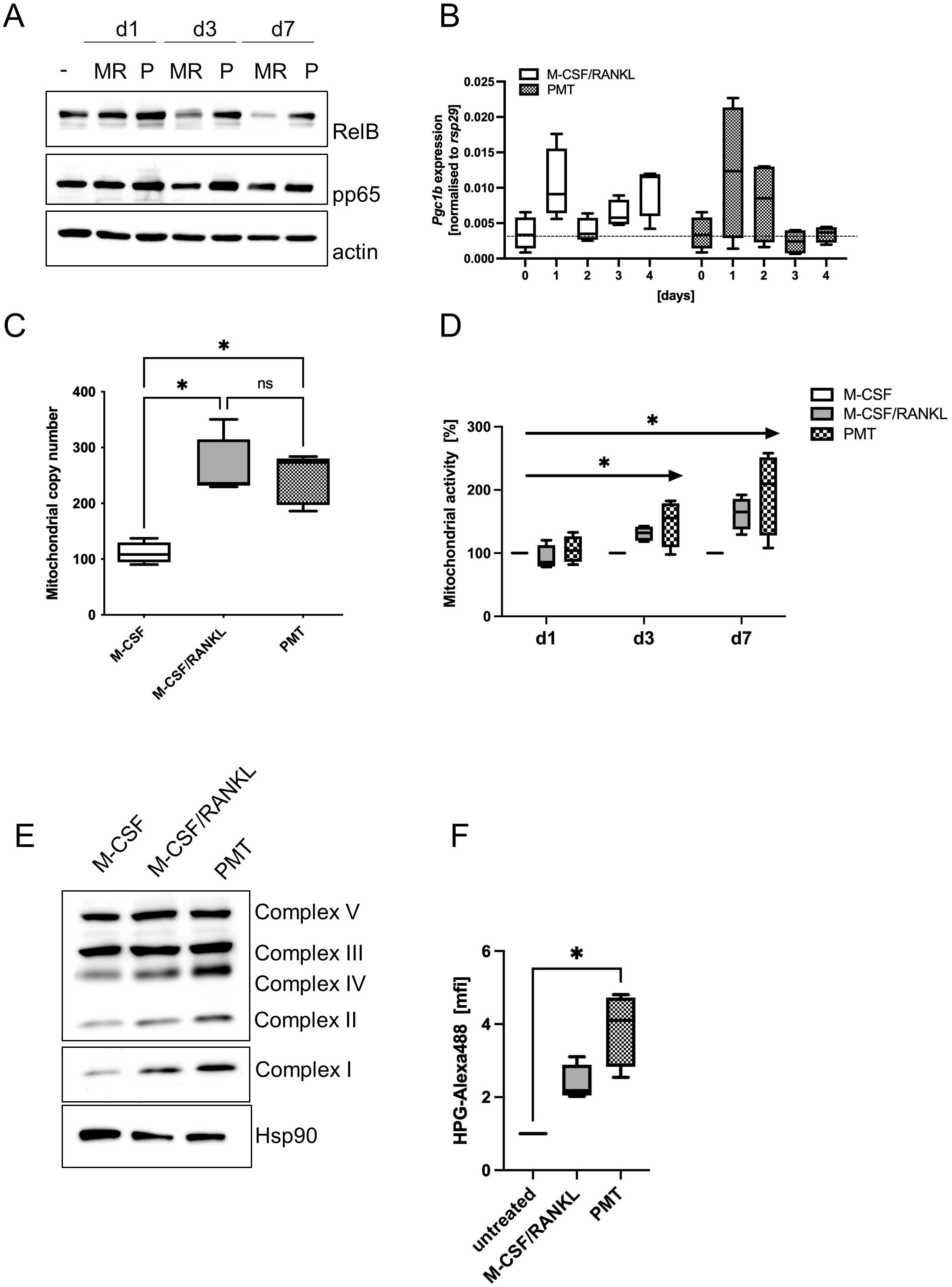
Mitochondrial activity during osteoclastogenesis. (A) BMDM were stimulated as indicated and lysates from day 0, 1, 3 and 7 were probed for the expression of RelB and pp65, actin was used as a lysate control (n=3). (B) Expression of *Pgc1b* was investigated by RT-PCR and normalised to the house-keeping gene *rsp29* (n=4). (C) BMDM were differentiated into OCs for 4 days before preparing genomic DNA. Mitochondrial copy number was determined by RT-PCR comparing the expression of a mitochondria specific sequence with nuclear DNA (n=4). (D) To quantify mitochondrial activity, differentiating cells were incubated at the indicated days with Mitotracker Deep Red and fluorescence intensity was quantified and normalised to the M-CSF control (n=3). (E) Expression of OxPhos proteins on day 3 of differentiation. Hsp90 was used as control (n=3). (F) Analysis of cellular translational activity using a HPG assay, were the methionine analogue HPG is coupled to Alexa488 on newly synthesised proteins (n=4). Statistical analysis was done using 2-way ANOVA for (B) and (C) and a Friedman test for (F) on raw data before normalisation.

### Gαq is abundantly expressed in the mitochondria of murine macrophages

To understand how PMT supports increased mitochondrial activity, we focussed on the PMT target Gαq itself. Overexpression of Gαq at the mitochondria had been shown to increase ATP production and to improve folding of the mitochondrial cristae in cardiomyocytes (Beninca et al., 2014). Therefore, we examined the localisation and activation of Gαq by PMT and found a preferential mitochondrial localisation and its deamidation at the mitochondria (Fig 3A). Beninca et al. suggested that Gαq supported mitochondrial activity by increasing the stability of the dynamin like GTPase OPA1, the main factor of cristae formation, which was supported by further studies using Opa1-deficient MEFs (Del Dotto et al., 2017). Because STAT3 is a central player in PMT-mediated signalling (Orth et al., 2007) and *Opa1* has a STAT3 binding site in its promoter (Nan et al., 2017), we investigated the role of STAT3-mediated transcription but did not detect major differences for *Opa1* between M-CSF/RANKL and PMT-treated cells (supp Fig 2A). To see if activation of mitochondrial Gαq by PMT would result in enhanced cristae folding, we investigated the expression of OPA1 after 72 h by western blotting. Indeed, OPA1 expression was significantly enhanced after PMT and M-CSF/RANKL addition (Fig 3B), which correlated with an increase in pSerSTAT3 for PMT-treated samples. As the expression of ETC proteins depends on the ability of the mitochondrial membrane to form extensive cristae (Beninca *et al*., 2014), this gives an additional explanation for the enhanced expression of ETC complexes after PMT treatment (Fig 2E). In addition to its role as a transcription factor, STAT3 also has non-genomic functions and can localise in mitochondria. Thus, we investigated if PMT treatment would also result in mitochondrial translocation of STAT3 and if that had an impact on OPA1 expression. As a control, we used the pro-inflammatory and osteoclastogenic STAT3-activating cytokine IL-6. Indeed, we found serine-phosphorylated STAT3 in the mitochondria after PMT and IL-6 treatment (Fig 3C).

**Figure 3:**
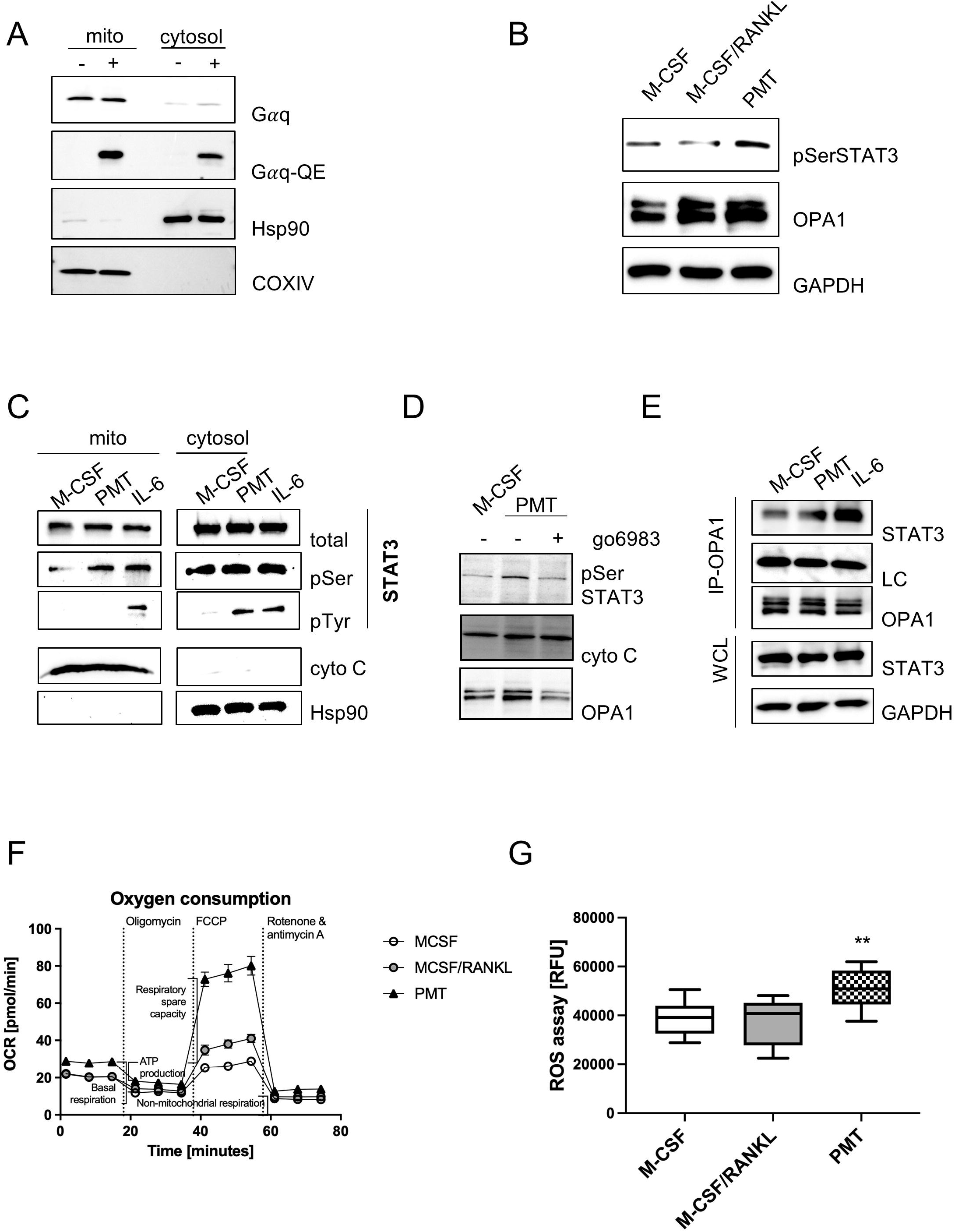
Gαq mediates STAT3 mitochondrial localisation and OPA1 expression. (A) Biochemical fractionation and subsequent analysis of mitochondrial and cytoplasmic fractions for deamidated and total levels of Gαq and OPA1, respectively (n=4). Hsp90 and COXIV were used as purity controls for cytoplasmic and mitochondrial fractions. (B) Whole cell lysates showing the serine phosphorylation of STAT3 and expression Opa1 on day3 of differentiation. (C) Localisation and post-translational modifications of STAT3 in mitochondrial and cytoplasmic fractions (n=4). (D) Serine phosphorylation of STAT3 and OPA1 expression were determined in mitochondrial fractions in the presence and absence of the pan-PKC inhibitor go6983 (n=3). (E) Whole cell lysates of BMDM were prepared after 3 h of stimulation with IL-6 or PMT and either used directly as WCL control or subjected to immunoprecipitation with a monoclonal OPA1 antibody. Membranes were immunoblotted using an antibody recognising total STAT3. As input control, a light chain specific antibody against the IP antibody was used. Equal OPA1 levels were verified after stripping the membrane. WCL samples were blotted against STAT3 and GAPDH as loading controls (n=4). (F) Seahorse analysis of mitochondrial activity on day3 of differentiation (Mito Stress Test) (n=3). (G) Quantification of ROS production of BMDM on day 3 of osteoclast differentiation (n=5, with triplicates). Statistical analysis was done using a Kruskal-Wallis test on unmatched samples.

Protein kinase C (PKC) is activated downstream of Gαq and has been reported to be able to phosphorylate STAT3 (Gartsbein et al., 2006). To see if Gαq-mediated PKC activation might trigger STAT3 serine phosphorylation and translocation, we inhibited PKC with the inhibitor go6983. This resulted in decreased pSer-STAT3 levels, as well as a substantial decrease in OPA1 expression (Fig 3D). As there was no difference in the Opa gene induction between RANKL and PMT samples, and only PMT caused mitochondrial STAT3 translocation, we hypothesised that STAT3 might stabilise OPA1 expression through a protein interaction. To this end, we performed co-precipitations using an OPA1 antibody and subsequent detection of precipitated STAT3 by immunoblotting. Again, IL-6 was used as a control. Figure 3E shows that PMT as well as IL-6 trigger the formation of an interaction between OPA1 and STAT3. This suggests that the observed decrease in OPA1 expression after inhibition of PKC is due to the missing interaction between STAT3 and OPA1 in the mitochondria. As OPA1 enhances cristae structures, we investigated if PMT could change the shape of mitochondria in H9c2-derived cardiomyocytes, as these cells are particularly rich in mitochondria. EM pictures indeed suggest that PMT causes a more refined structure of cristae (supp. Fig 2B).

Increased OPA1 protein expression was described to enhance complex I and II activity and the oxygen consumption rate (OCR) and the production of ATP (Nan *et al*., 2017; Rincon and Pereira, 2018), therefore, we investigated if the Gαq-mediated effects on cristae structure and OxPhos expression resulted in a change in OCR. Mitochondrial activity was measured using a Seahorse mito stress test assay. As expected, both types of OCs showed an altered mitochondrial respiration when compared with macrophages. However, the respiratory indices of PMT-treated OCs were much higher than those of cytokine-induced OCs (Fig 3F, supp. Fig 2C-E). As both osteoclasts have increase OPA1 expression, the additional interaction withSTAT3 which happens only in PMT-treated cells, seems to be central for the additional increase in OCR. As a consequence, PMT-treated cells also produce higher amounts of ROS (Fig 3G), which is a secondary messenger for osteoclastogenesis and should thus support cell differentiation. In addition, oxidative stress was not increased in PMT-treated cells, as they showed an increase in glutathione levels (supp Fig 2F).

### Gαq overexpression causes a phenotype of hypermetabolic activity

To prove that the effects observed for treatment with PMT are actually driven by the constitutive activation of Gαq signalling, we retrovirally overexpressed Gαq in ER-Hoxb8 cells (supp Fig 3A). Figure 4A shows successful transduction by verifying enhanced Gαq expression levels. TRAP assays were performed and the number of differentiated cells was determined. Figure 4B,C shows that Gαq overexpression increased the differentiation into TRAP-positive, multinucleated cells. The higher variability observed for the number of TRAP-positive, multinucleated ER-Hoxb8-Gαq cells could be caused by the fact that Gαq cells were slower in adhering to the wells, thus having a slower kinetics and a higher number of yet smaller-sized cells.

**Figure 4:**
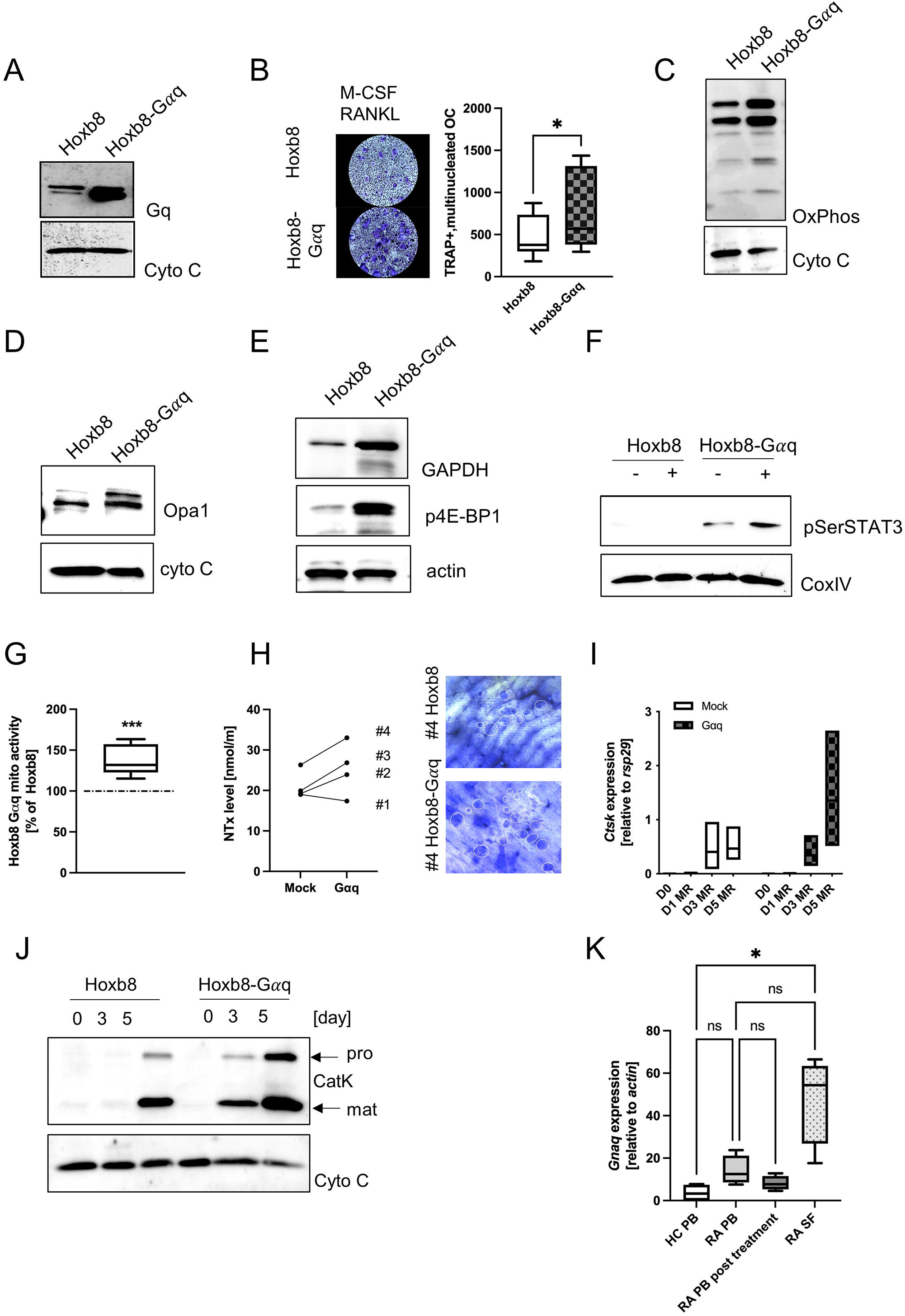
Overexpression of Gαq in ER-Hoxb8 cells causes a hypermetabolic phenotype. (A) ER-Hoxb8 cells were retrovirally transduced with murine Gαq by transduction and overexpression was verified by immunoblotting. Cytochrome c was used as lysate control. (B) TRAP assays were performed to study the effect of Gαq on OC formation. Cells were stained for TRAP and multinucleated, TRAP positive cells were quantified (n=4). Lysates were probed for the expression of OxPhos components (C), OPA1 (D), GAPDH and p4E-BP1 (E). Cytochrome C and actin were used as controls (n=3). (F) ER-Hoxb8 and ER-Hoxb8-Gαq cells were left untreated or stimulated with PMT for 3 hours before lysis. Lysates were probed for pSer STAT3; COXIV was used as a lysate control (n=3). (G) Mitochondrial activity was quantified by Mitotracker Deep Red analysis and subsequent evaluation of mfi values (n=5). Statistical analysis was done performing a Wilcoxon test on the raw data before normalisation to ER-Hoxb8 as 100%. (H) Bone slices were incubated for 14 days with ER-Hoxb8 and ER-Hoxb8-Gαq cells before staining the bone with toluidine blue. Supernatants were used to quantify the released collagen (day3) (n=4). Statistical analysis was done using a Wilcoxon test. The bone slices shown correspond to #4 of the ELISA. (I) RT-PCR analysis of ER-Hoxb8 and ER-Hoxb8-Gαq for *Ctsk* induction normalised to *rsp29* on days 0, 1, 3 and 5 (n=3). Statistical significance was evaluated with a 2-way ANOVA test. (J) Cathepsin K Western Blot with lysates from day 0 - d3 of ER-Hoxb8 and ER-Hoxb8-Gαq. (n=3). (K) *Gnaq* expression was quantified by RT-PCR using peripheral blood from healthy controls and RA patients, as well as synovial fluid from RA patients. In addition, PB of RA patients post-treatment was analysed (n=4). Statistical analysis was done using a Friedman test.

To see whether Gαq overexpression would be sufficient to mimic the phenotype of PMT-induced constitutive Gαq activation, we checked the expression pattern of central signalling molecules. Indeed, we could corroborate an increase in OxPhos expression (Fig 4C), OPA1 (Fig 4D), GAPDH as well as p4-EBP1 (Fig 4E). Also, we observed an increase in pSer STAT3 levels that was caused by overexpression of Gαq and was enhanced by PMT treatment (Fig 4F). As expected, we could also observe higher mitochondrial activity (Fig 4G). To see if the enhanced metabolic activity would result in increased resorptive activity, we performed RT-PCR analysis for activity-related genes (*Ctsk, Mmp9, Acp5* and *Dcstamp*), western blot analysis for Cathepsin K expression and cleavage into its active form as well as bone resorption assays and subsequent quantification of released collagen (Fig 4H-J and supp Fig 3B-D). Again, Gαq cells were more active than the control and the enhanced expression of Cathepsin K on gene and protein level relative to the corresponding control suggests that this not only caused by an accelerated differentiation but also by enhanced functional activity.

Therefore, we asked whether an increase in Gαq expression might also be observed in patients with increased osteoclastogenesis such as rheumatoid arthritis. For this pathology, it was described that patient-derived macrophages display a phenotype of hyperactivated metabolism (Weyand et al., 2017). When we investigated the expression of *Gnaq* in the peripheral blood from healthy donors and RA patients, we observed a trend towards increased *Gnaq* in RA patients compared to the healthy control. This increase in *Gnaq* was even higher when synovial fluid samples were used for analysis (Fig 4K). Interestingly, PB from RA patients post-treatment showed decreased levels of *Gnaq* and *Opa1* gene induction, suggesting that both factors are central players for disease severity (supp Fig 3E).

## Discussion

OC have to adapt their metabolic activity throughout their differentiation process to enable cell fusion and resorption of inorganic material (Kubatzky et al., 2018). Our data show that osteoclasts differentiating with PMT display a distinct phenotype where glycolysis, mTOR pathway and translational activity are strongly activated. As a consequence, expression of OxPhos proteins and mitochondrial activity is elevated. Gαq-dependent phosphorylation of STAT3 is a central step as it promotes mitochondrial translocation and increases mitochondrial activity. Consequently, overexpression of Gαq in ER-Hoxb8 cells was sufficient to mimic this phenotype. Our data suggest that mitoSTAT3 helps restructuring mitochondria through increased expression or stability of OPA1, a GTPase required for efficient cristae folding. Through this, OPA1 increases mitochondrial respiration and helps to maintain functionality of the TCA cycle (Cogliati et al., 2016). Correspondingly, in Gαq/11-deficient fibroblasts, OPA1 levels as well as the numbers of cristae were reduced, causing narrow junctions and reduced ATP synthesis (Beninca *et al*., 2014). MitoSTAT3 was described to interact with ETC complexes to support electron flux and increase mitochondrial metabolism (Garama et al., 2016). In cardiomyocytes, mitoSTAT3 enhanced OxPhos activity and acted protectively during ischaemia/reperfusion (Yang and Rincon, 2016) and in cancer this promotes reprogramming from glycolysis to OxPhos (Lee et al., 2019). Our data suggest that this may additionally be supported by OPA1, another well-known mediator of cardiomyocyte resistance to apoptosis. The observed increase in OPA1 expression seems to be mediated through the direct association with mitoSTAT3 as inhibition of STAT3 translocation to the mitochondria, decreases Opa1 expression. This would represent a novel function for mitoSTAT3 that was previously shown to act as a stabilising factor in the mitochondria through its interaction with GRIM19 of the ETC complex I (Huang et al., 2004; Wegrzyn et al., 2009). Little is known about a possible function of mitoSTAT3 in immune cell regulation. Rincon et al. suggest that IL-21 and IL-6 play a role in mitochondrial functions of STAT3 in lymphocytes. In CD4 T cells, IL-6 causes mitochondrial hyper-polarization and increases mitochondrial calcium. As a consequence, cytosolic calcium levels increase and support persistent cytokine expression (Rincon and Pereira, 2018; Yang et al., 2015). The cytokine RANKL does not activate STAT3 suggesting that mitoSTAT3 is not involved in OC formation under physiological conditions. However, in an inflammatory microenvironment, IL-6 is highly expressed by macrophages and potentially triggers STAT3 serine phosphorylation as well as its mitochondrial translocation. In RA patient lymphocytes, enhanced nuclear STAT3 activity was observed and inhibition of STAT3 tyrosine phosphorylation or overexpression of the mitoSTAT3 interaction partner Grmp19 ameliorated RA as it reduced excess glycolytic activity, increased OxPhos activity and reduced the amount of OC stimulating Th17 T cells (McGarry et al., 2018; Moon et al., 2014).

Macrophages from RA patients display a distinct, hypermetabolic phenotype, characterised by the dual activation of glycolysis and OxPhos (Weyand *et al*., 2017; Zeisbrich et al., 2018). As we had observed that overexpression of Gαq causes a similar hypermetabolic state, we wanted to know if Gαq overexpression might be of pathophysiological relevance in RA. Indeed, we found *Gnaq* to be upregulated in peripheral blood and synovial fluid from RA patients. This is in contrast with other studies that found a lower *Gnaq* expression in RA lymphocytes (Wang et al., 2012). Interestingly, there is also a gender bias as the expression of Gnaq increases in an oestrogen-dependent manner in immune cells (Morton et al., 2003). In addition, as outlined above, the metabolic regulation of lymphocytes and macrophages from RA patients is fundamentally different. It is therefore possible that Gαq plays a role in the mitochondrial hyperactivity observed in macrophages but is decreased in lymphocytes for the same reason.

It has been suggested that improving mitochondrial physiology might be an interesting option to reduce inflammation in RA (Castegna et al., 2020), but it can be anticipated that immunometabolic targets need to be addressed in a cell type specific manner. Gαq was reported to be down-regulated in RA patient lymphocytes (Liu et al., 2015). Targeting Gαq in macrophages might therefore be an interesting option that circumvents the dual roles observed for other proteins, such as mTOR, STAT3 or HIF-1α. It is currently discussed to target G protein-coupled receptors (GPCR) signalling by developing inhibitors for their downstream subunits (Campbell and Smrcka, 2018; Kostenis et al., 2020). This has the advantage that receptors that employ more than one heterotrimeric G protein can be targeted pathway specifically. GPCRs play critical roles in a number of clinically relevant diseases, such as cancer, cardiovascular diseases as well as infection and inflammation (Jo and Jung, 2016). Chemokine receptors for example determine the spatio-temporal organisation of the leukocyte immune response (Lammermann and Kastenmuller, 2019). Not surprisingly, these receptors have also been implicated in the pathogenesis of autoimmune pathologies like SLE, rheumatoid arthritis and systemic sclerosis, where GPCR activation could be linked to Gαq activation downstream in same cases (Zhang and Shi, 2016). More importantly, GPR91, the receptor for the TCA metabolite succinate, was found to signal via Gαq in macrophages (Trauelsen et al., 2021). This GPCR can activate pro- and anti-inflammatory signalling as well osteoclastic gene expression via Gαq-mediated activation of NFATc1 (Guo et al., 2017). GPR91 represents a central molecular switch in RA and correspondingly inhibition or deletion of GPR91 ameliorated the disease (Littlewood-Evans *et al*., 2016; Saraiva et al., 2018; Velcicky et al., 2020). For Gαq, two bioavailable inhibitors exist at the moment (YM254890, FR900359) (Schrage et al., 2015; Taniguchi et al., 2003). However, only preclinical studies for their effectiveness in melanoma, asthma and thrombosis have been performed to date and the suitability for treatment of autoimmune diseases remains to be investigated. Our data suggest that inhibition of Gαq might be an interesting drug target to modulate cellular metabolic activity.

## Materials and Methods

### Mice and BMDMs culture

6- to 8-week-old female C57BL/6J mice were bred under SPF conditions and sacrificed in accordance with the animal care guideline approved by German animal welfare authorities. Bone marrow cells were isolated from mice femur and tibia, and gently washed with 1x PBS. Cells were maintained in high glucose Dulbecco’s medium supplemented with 10% FCS, 1% penicillin/streptomycin, and 50 µM ß-mercaptoethanol (DMEMs) plus the source of M-CSF derived from L929 cell lines. Cells cultured for 3 days were re-stimulated with additional L929-derived M-CSF to be differentiated into bone marrow-derived macrophages (BMDMs). Primary macrophages were harvested on day 6 and were stimulated with 25 ng/ml M-CSF, 50 ng/ml RANKL, 50 ng/ml IL-6 or 1 nM PMT as indicated in figure legends.

### Retroviral transduction of Hoxb8 cells

pMX-Gαq-IRES-CD4 or pMX-IRES-CD4 (control) was transiently transfected into phoenix-eco cells using ProFection® Mammalian Transfection System. Experimental procedures were performed according to the manufacturer’s instructions. About 4 to 8 × 10^5^ phoenix-eco cells were seeded per well of a 6-well plate in Dulbecco’s medium supplemented with 10% FCS and 1% penicillin/streptomycin a day prior to transfection. Retroviral supernatant was collected 2 days post transfection and used for infection of ER-Hoxb8 conditionally immortalized murine macrophage progenitors. About 5 × 10^5^ Hoxb8 cells in virus supernatant along with 16 ug polybrene were centrifuged at 1100 rpm (37°C) for 2 hours. Cells were further cultured in RPMI 1640 medium with 10% FCS, 1% penicillin/streptomycin, 5% GM-CSF supernatants, 1 µM β-oestradiol. Additional 2.5 g/L glucose was supplemented in RPMI 1640 medium for efficient cell viability. Infected Hoxb8 cells harbouring DNA constructs were selected with 8 ug/ml puromycin. After 6 days of incubation, transduction efficiency was subsequently confirmed by CD4 expression based on flowcytometry analysis. Culture media was completely refreshed every 2 days. Stable Hoxb8 cell lines were gently washed twice in RPMI1640 medium to remove β-estradiol, which blocks cellular differentiation. Ready-to-use Hoxb8 cells were cultured in DMEMs for further experiments.

### Osteoclast differentiation and TRAP staining

7 × 10^4^ Hoxb8 cells were seeded in 48-well plates. Cells were stimulated with M-CSF and RANKL; after every 3 days half of the culture medium was replenished with medium containing M-CSF and RANKL. Multi-nucleated cells were observed under a light microscope Axiovert 25 typically after 4 to 5 days in culture. Cells were fixed and stained using the Leukocyte Acid Phosphatase (TRAP) kit according to the manufacture’s recommendations. TRAP-positive cells with more than 3 nuclei were counted as mature osteoclasts.

### Bone resorption assay

2 × 10^5^ Hoxb8 or Hoxb8-Gαq cells were seeded per well in a 24-well plate and stimulated with indicated M-CSF and RANKL for 3 days. For the bone resorption assay, Cortical bovine bone slices were used. Cortical bovine bone slices were washed with medium several times in order to remove the alcohol and placed in the 96 well plate. After 3 days of stimulation, cells were gently detached by Accutase solution and transferred onto bone slice in a 96-well plate. Cells were kept on bone slices for 15 days; every third day, half of the medium was replenished with medium containing MCSF and RANKL. After thoroughly removing the cells from the bone slices, using freshly prepared 0.1% toluidine blue, pits were stained on the bone slices. Pit formation on bone slices were analysed with a Rebel Microscope (ECHO A Bico Company).

### NTX ELISA

Cell supernatants collected on day 9 during bone resorption assay were used to determine levels of the secreted N-telopeptide of type 1 collagen. Experimental details were followed by manufacture’s protocol. Cell supernatants were added along with NTX biotinylated antibody and Streptavidin-HRP into a pre-coated well with NTX antibody for 1 hour at 37°C. After 1 hour of incubation, wells were washed 5 times with wash buffer. The substrate solution was added to develop the colour and the reaction was stopped using stop solution. The absorbance was measured at 450 nm using a CLARIOstar (BMG Labtech).

### Real-time PCR

Total RNA was isolated from cells using innuPREP RNA Mini Kit 2.0. RNA isolation was performed in accordance with the provided manual and quantified by a Nanodrop. A concentration of 500 ng/ul of total RNA was converted to cDNA using cDNA synthesis kit. Real-time PCR analysis was further performed on a StepOnePlus Real-Time PCR System from Qiagen using 2x qPCRBIO SyGreen Mix Hi-ROX. Specific primer pairs can be found in table 1. Relative gene expression of target genes was calculated in comparison to Ct value of house-keeping gene of Rsp29 using 2^-[Ct(target gene)−Ct(reference gene)]^.

### Expression of GNAQ and OPA1 in human samples

The study was conducted with approval from Institute Ethics Committee (IEC-490/01.09.2017). After obtaining informed consents, patients with RA were recruited from Orthopaedics OPD of AIIMS, New Delhi [Age in years (Mean ± SD): 41±11 years; Sex: Female]. Diagnosis was made on the basis of ACR criteria 1987. For this study, we collected synovial fluid (SF) and autologous peripheral blood (PB) from RA patients with active disease. We also collected peripheral blood of RA patients with low disease condition (Under treatment) [Age in years (Mean ± SD): 45±6 years; Sex: Female]. Healthy controls (HC) collected for the studies, were free from any acute or chronic ailment and were not on medication at the time of enrolment. Peripheral blood from HC was collected [Age in years (Mean ± SD): 35±5 years; Sex: Female]. Specimens were collected in heparinized tubes.

Synovial Fluid mononuclear cells (SFMC) and Peripheral blood mononuclear cells (PBMCs) were isolated using Lymphoprep, density gradient centrifugation. Cells were resuspended in RPMI-1640 containing L-Glutamine, and HEPES supplemented with, 10% FBS, 100 U/ml penicillin and 100 µg/ml streptomycin. A total of 5 × 10^6^ cells were seeded per well of 6 well plate. Cells were kept in humidified 5% CO_2_ incubator at 37 °C for 24 hours. After 24 hours, non-adherent cells were removed by washing the wells with PBS three times. Adherent cells were processed for expression studies. RNA was extracted using GeneJET RNA Purification Kit, according to the manufacturers’ protocol. cDNA was prepared by using Revert Aid First strand cDNA synthesis kit. Quantitative RT-PCR was performed using Powerup Sybr Master Mix with the primers mentioned. RT-PCR was performed using the QuantStudio 5 Real-Time PCR Systems (Applied Biosystems). An initial enzyme activation step of 2 minutes at 50°C and denaturation step of 5 minutes at 95°C, followed by amplification for 40 cycles at 95°C for 15 seconds and at 56°C for 1 minute. As normalization control ACTB, was used. Relative gene expression of target genes was calculated in comparison to Ct value of normalizing control using 2^-[Ct(target gene)−Ct(reference gene)]^.

### Immunoblotting analysis

Cells were lysed using a RIPA buffer (1% NP-40, 0.25% deoxycholate, 50 mM Tris pH7.4, 150 mM NaCl, 1 mM EDTA, 1 mM Na_3_Vo_4_) supplemented with a Phosphatase and Protease-Inhibitor Cocktail. Cell lysates were obtained after shaking for 45 minutes in cold room, followed by centrifugation for 20 minutes at 15,000 rpm at 4°C. About 10 ug of proteins were subjected to ProGel Tris Glycine 4-20% polyacrylamide gel, and then transferred to the nitrocellulose blotting membrane via semi-dry blot system. Membranes were blocked in 1x blocking buffer for 30 minutes at room temperature and incubated with the desired primary antibody overnight at 4°C. Probed bands were detected by the appropriate HRP-conjugated secondary antibody upon incubation for 1 hour at room temperature. Membranes were developed using the ECL substrate WESTAR *η*C Ultra 2.0 and visualized on ChemoStar image (Intas Science Imaging).

### Co-immunoprecipitation

1.5 × 10^7^ stimulated BMDMs were lysed in 1x Brij buffer with 1% Brij97, 150 mM NaCl, 20 mM Tris pH7.4, 1 mM EDTA, 1 mM MgCl_2_, 1mM Na_3_VO_4_, and 10% Glycerol with a Phosphatase and Protease-Inhibitor Cocktail. Total lysates were obtained as described above. Collected samples were incubated with protein A/G plus agarose beads and anti-mouse Opa1 antibody overnight at 4°C. Beads were gently washed twice with Brij buffer and once with TNE buffer for 15 seconds and used for immunoblotting.

### Mitochondrial fractionation

Biochemical mitochondria fractionation was performed based on the sequential centrifugation using Mitochondrial Isolation Kit for Mammalian Cells according to manufacturer’s instructions with a minor modification. 2 × 10^7^ of BMDMs were lysed with isolation reagents supplemented with 20x Protease inhibitor. Samples were homogenized with 30 strokes using a glass Dounce tissue grinder. Every centrifugation step was conducted at 4°C. Homogenates were centrifuged for 10 minutes at 3000 rpm and supernatants were transferred for additional centrifugation for 15 minutes at 12000 rpm. Supernatants containing cytosolic proteins were obtained in this step. The pellet was washed for 5 minutes at 12000 rpm and resuspended in 2% CHAPS buffer with 25 mM Tris-HCl, 0.15 M NaCl, pH7.2. Mitochondrial fractions as supernatants were isolated after centrifugation for 2 minutes at 13000 rpm.

### Mitochondrial DNA copy number

Total DNA was extracted using DNAeasy Blood and Tissue Kit following the provided protocol. About 10 ng of DNA was amplified by RT-PCR as described above. Relative mitochondrial DNA contents were normalized to the level of nuclear DNA (see primer sequences in table 1). Relative mitochondrial DNA copy was calculated as 2×2^(nucleus Ct-mitochondria Ct)^.

### Translation assay

Translational activity was achieved by the methionine analogue of L-homopropargylglycine (HPG) incorporation via Click-iT® HPG Alexa Fluor® 488 Protein Synthesis Assay Kit. 1 × 10^6^ of BMDMs in a 24-well plate were stimulated for 6 hours, and 50 µM HPG was added 30 minutes before stimulation was over. Samples were fixed in 4% PFA and permeabilized with 0.5% Saponin. Click-iT® reaction cocktail including Alexa Fluor 488 dye was added in each well for 40 minutes at room temperature under dark condition. Protein synthesis was evaluated by measuring FITC channel for Alexa Fluor 488 on BD FACSDiva software using a BD FACSCanto Cytometer.

### Seahorse assay

Life cell metabolic analyses were performed measuring extracellular acidification rate (ECAR) and oxygen consumption rate (OCR) using Seahorse XFp Analyzer (Agilent Technologies). 4×10^5^ cells per well were seeded and incubated for 3 days. One hour before the measurements, the growth medium was replaced by Seahorse XF Base Medium (without Phenol Red) supplemented with 5 mM HEPES, 1mM sodium pyruvate, 2 mM L-glutamine, only for assays analysing mitochondrial respiration, 10 mM D(+)Glucose. Thereafter, cells were incubated at 37°C without CO_2_ until start of the assay. After three baseline measurements (each measuring point comprises 3 min mixing and 3 min measuring) 2 µM oligomycin, 1.5 µM Carbonyl cyanide–4 (trifluoromethoxy) phenylhydrazone (FCCP), and 0.5 µM rotenone/antimycin A (all three from Seahorse XFp Cell Mito Stress Test Kit) were added sequentially to characterize mitochondrial respiration. To determine glucose uptake and glycolytic function 10mM glucose, 2µM oligomycin, and 50 mM 2–Deoxy-D–glucose (2-DG) (all three from Seahorse XFp Glycolysis Stress Test Kit) were added sequentially. OCR and PER were calculated using the Wave 2.6.0 software and parameters were calculated according to the manufacturer’s instruction using Seahorse XF Cell Mito Stress Test Report Generator 3.0.11 and Seahorse XF Glycolysis Stress Test Report Generator 4.0.

### Mitochondrial activity

For FACs analysis, 1 × 10^6^ cells were used per sample. Mitochondrial activity was determined by staining of cells for 30 minutes with 100 nM MitoTracker Deep Red. Cells were then washed three times with 1x PBS and the mean fluorescence intensity (MFI) was measured in the APC-Cy7 channel.

### 2-DIGE

A difference-in-gel electrophoresis (DIGE) technique was used to investigate proteome differences between PMT-stimulated and M-CSF/RANKL and M-CSF-stimulated cells. 3×10^7^ cells were seeded to generate osteoclasts from primary macrophages. Cells were washed before lysis in 150 µl THC buffer. 100 μg of protein samples were stained using the Amersham CyDye DIGE Fluor minimal dye labelling kit according to the manufacturer’s instructions. All samples were stained once with Cy5 and once with Cy3, while the normalization control (all samples combined) was stained with Cy2. The samples to be compared (Cy2 and Cy3 stained) were then mixed with the Cy2-stained normalization control and separated two-dimensionally according to isoelectric point and molecular weight. The samples were then applied to IPG strips and separated in the first dimension with the IPGphor 3 system from GE Healthcare according to their isoelectric focusing (20 C, I < 25 μA). After equilibration of the IPG strips in reduction buffer and alkylation buffer proteins were separated according to their molecular weight in the second dimension using a gradient gel (8%-15%) in an Ettan DALT II system (GE Healthcare). The gels were then scanned at the different wavelengths using a Typhoon 9400 Laser Scanner (GE Healthcare) and analysed using the DeCyder 2D software. The spots were automatically picked with an Ettan Spot Picker (GE Healthcare) and transferred to a 96-well plate. The spots were analysed by mass spectrometry (LTQ Orbitrap XL mass, Thermo Fisher Scientific) by Dr. Martina Schnölzer (DKFZ, Heidelberg).

### Total ROS assay

5×10^4^ cells were seeded in black 96-well plates and stimulated as indicated. On day 3 the cells were processed according to the manufacturer’s protocol. Fluorescence at 520 nm was measured using a FluoStar Optima (BMG Labtech, Offenburg, Germany).

### GSH/GSSG-Glo Assay

For this assay (Promega, Germany), 2×10^4^ cells were seeded in a 96-well plate and stimulated as indicated. On day 3 the cells were processed according to the manufacturer’s protocol. Luminescence was measured in white 96-well plate using a LumiStar Optima (BMG Labtech, Offenburg, Germany). The concentration of GSH was calculated using a standard curve with glutathione (16 μM to 0.125 μM).

### Data analysis

All data presented as mean ± SD. Statistical analyses were carried out using GraphPad Prism 9.0 software as indicated in the respective figure legend.

| REAGENT or RESOURCE | SOURCE | IDENTIFIER |
| --- | --- | --- |
| Antibodies |  |  |
| Rabbit monoclonal anti-pAMPKα (Thr172) | Cell Signaling | 2531 |
| Mouse monoclonal anti-GAPDH | Proteintech | 60004-1-Ig |
| Rabbit polyclonal anti-pGSK-3β (Ser9) | Cell Signaling | 5558 |
| Rabbit monoclonal anti-β-Actin | Proteintech | 20536-1-AP |
| Rabbit polyclonal anti-p-Zap-70 (Tyr319)/Syk (Tyr352) | Cell Signaling | 2701 |
| Rabbit monoclonal anti-pAkt (Thr308) | Cell Signaling | 2965 |
| Rabbit monoclonal anti-p4E-BP1 (Thr37/46) | Cell Signaling | 2855 |
| Rabbit monoclonal anti-pmTOR (Ser2448) | Cell Signaling | 5536 |
| Rabbit monoclonal anti-Relb | Cell Signaling | 4922 |
| Rabbit monoclonal anti-pNF-κB p65 (Ser536) | Cell Signaling | 3033 |
| Rabbit monoclonal anti-pNF-κB p105 (Ser932) | Cell Signaling | 4806 |
| Mouse monoclonal anti-total OXPHOS cocktail | Abcam | Ab110413 |
| Rabbit polyclonal monoclonal anti-Hsp90 | Cell Signaling | 4874 |
| Mouse monoclonal anti-Gαq | Santa Cruz | sc-136181 |
| Rabbit monoclonal anti-CoxIV | Cell Signaling | 4850 |
| Rat anti-Gq(QE) | Kamitani et al., 2011 | N/A |
| Rabbit monoclonal anti-OPA1 | Cell Signaling | 80471 |
| Rabbit monoclonal anti-pStat3 (Ser727) | Cell Signaling | 94994 |
| Rabbit monoclonal anti-pStat3 (Tyr705) | Cell Signaling | 9145 |
| Rabbit monoclonal anti-STAT3 | Cell Signaling | 4904 |
| Rabbit polyclonal anti-Cytochrome c | Proteintech | 10993-1-AP |
| Goat polyclonal anti-mouse IgG light chains | Jackson/ Dianova | 115-035-174 |
| Goat monoclonal anti-mouse IgG, HRP-linked | Cell Signaling | 7076 |
| Goat monoclonal anti-Rabbit IgG, HRP-linked | Cell Signaling | 7074 |
| Mouse anti-human CD4 APC | Miltenyi Biotec | 130-113-210 |
| Biological Samples |  |  |
| Peripheral blood and Synovial fluid of Rheumatoid arthritis; Peripheral blood of Healthy subjects | All India Institute Of Medical Science, New Delhi | Institute Ethics Committee (IEC-490/01.09.2017) |
| Chemicals, peptides, and recombinant proteins |  |  |
| MitoTracker® Deep Red FM | Cell Signaling | 8778 |
| Gö 6983 | Merck | 365251 |
| Pasteurella multocida toxin (PMT) | Chakraborty et al., 2017 | N/A |
| Human IL-6 | Miltenyi Biotec | N/A |
| Recombinant Mouse M-CSF Protein | Biotechne | 416-ML |
| Recombinant Mouse TRANCE/RANK L/TNFSF11 Protein | Biotechne | 462-TR |
| CellROX™ Deep Red Reagent | Thermo Fisher | C10422 |
| Puromycin | Sigma-Aldrich | P8833 |
| Protein A/G plus agarose | Santa Cruz | sc-2003 |
| β-estradiol | Sigma-Aldrich | E2758 |
| β-mercaptoethanol | PAN Biotech | P07-05020 |
| Critical commercial assays |  |  |
| Click-iT™ HPG Alexa Fluor™ 488 Protein Synthesis Assay Kit | Thermo Scientific | C10428 |
| QPRO-BCA Kit Standard | Cyanagen Srl | PRTD1,0500 |
| Total ROS Assay Kit 520 nm | Thermo Scientific™ | 88-5930-74 |
| innuPREP RNA Mini Kit 2.0 | Analytikjena | 845-KS-2040250 |
| GeneJET RNA Purification Kit | Thermo Scientific™ | K0732 |
| Revert Aid First strand cDNA synthesis kit | Thermo Scientific™ | K1621 |
| PowerUp™ SYBR™ Green Master Mix | Applied Biosystems™ | A25741 |
| cDNA synthesis kit | Biozym Scientific | 331470L |
| 2x qPCRBIO SyGreen Mix Hi-ROX | PCR Biosystems | PB20.12-01 |
| DNAeasy Blood and Tissue Kit | Qiagen | 69504 |
| Mitochondria Isolation Kit for Mammalian cells | Thermo Fisher | 89874 |
| ProGel Tris/Glycin gel 4-20% 1.0 mm | Anamed | TG42012 |
| WESTAR ηC Ultra 2.0 | Cyanagen Srl | XLS075,0100 |
| Leukocyte acid phosphatase (TRAP) Kit | Sigma-Aldrich | 387A-1KT |
| Bovine Cross linked N-telopeptide of type I collagen, NTX ELISA Kit | MYBioSource | MBS1602302 |
| Toluidine Blue O | Sigma-Aldrich | T3260 |
| Accutase Cell Detachment Solution | PAN Biotech | P10-21100 |
| D(+)-Glucose | Sigma-Aldrich | N/A |
| ProFection® Mammalian Transfection System | Promega | E1200 |
| GSH/GSSG-Glo™ Assay | Promega | V6611 |
| Amersham CyDye DIGE Fluor Minimal Labeling Kit | Cytiva | 25801065 |
| Sodium pyruvate | Thermo Fisher | N/A |
| L-glutamine | Thermo Fisher | N/A |
| Penicillin-Streptomycin Solution 100X | Anprotec | AC-AB-0024 |
| Seahorse XFp Cell Mito Stress Test Kit | Agilent | N/A |
| Seahorse XFp Cell Glycolysis Stress Test Kit | Agilent | N/A |
| Seahorse XF Base Medium | Agilent | N/A |
| HEPES | Agilent | N/A |
| Lymphoprep | Axis-Shield | 1114544 |
| FCS | Biochrome | N/A |
| RPMI 1640 w/ stable Glutamine | Anprotec | AC-LM-0056 |
| RPMI-1640 containing L-Glutamine, and HEPES | Gibco | 22400089 |
| DMEM High Glucose w/ stable Glutamine w/ Sodium Pyruvate | Anprotec | AC-LM-0013 |
| Experimental models: Cell lines |  |  |
| Mouse cell line: L-929 | ATCC | CCL-1 |
| Mouse cell line: ER-Hoxb8 | Wang et al., 2006 | N/A |
| Mouse cell line: LGM3 (muGM-CSF) | Zal et al., 1994 | N/A |
| Human cell line: Phoenix-Eco | ATCC | SD-3444 |
| Experimental models: Organisms/strains |  |  |
| Mouse: C57BL/6 | Janvier-labs | N/A |
| Oligonucleotides (Accession No.) |  |  |
| Recombinant DNA |  |  |
| pMX-IRES-CD4-Puro | Gündogdu et al., 2010 | N/A |
| pMX-Gq-IRES-CD4-Puro | This paper | N/A |
| pCl-eco | Naviaux et al., 1996 | N/A |
| Software and algorithms |  |  |
| FACSDiva™ | BD Biosciences | https://www.bdbiosciences.com/ |
| bioRENDER | bioRENDER | https://biorender.com/ |
| DeCyder 2D Version 7.0 | Amersham Bioscience | N/A |
| Flowing Software 2 | TURKU BIOSCIENCE | https://bioscience.fi/ |
| Fiji | Image J | https://imagej.net/software/fiji/ |
| Rebel Microscope | ECHO | https://discover-echo.com/ |
| PANTHER Classification System | PANTHER | http://www.pantherdb.org/index.jsp |
| Seahorse XF Cell Mito Stress Test Report Generator 3.0.11 | Agilent | N/A |
| Seahorse XF Glycolysis Stress Test Report Generator 4.0 | Agilent | N/A |
| GraphPad Prism 8 | GraphPad | https://www.graphpad.com/ |

## Acknowledgements

The study was founded partly through the priority program SPP1468 IMMUNOBONE by the Deutsche Forschungsgemeinschaft (DFG). S.C is supported by DBT Wellcome Trust India Alliance (IA/E/16/1/503016) Early career fellowship. We acknowledge Dagmar Schnölzer (DKFZ, Heidelberg) for performing Mass Spectrometry analysis, The EM facility (Heidelberg University), Gabriele Sonnenmoser for technical assistance, Elisabeth Seebach and Kathrin Thedieck for helpful discussions. We would like to thank the following students for their specific contributions: Lisa Rausch (DIGE), Lisa Baur (EM), Emma Pietsch (Glutathione Assays), David Reeg and Nicolas Stevens (Hoxb8 cells).

## Author Contributions

Conceptualization: K.F.K., K.B. and F.U.; Methodology, S.C., B.H., and K.B.; Investigation: S.C., B.H., D.Y., A.H., and J.S..; Formal Analysis: D.S.; Resources: T.T., S.K. and T.V.; Writing – Original Draft, K.F.K. and S.C.; Writing, Review & Editing: K.F.K., S.C. and B.H.; Funding Acquisition K.F.K. and S.C.

