## Supplementary Figures for "Gα_q_ modulates the energy metabolism of osteoclasts"

A

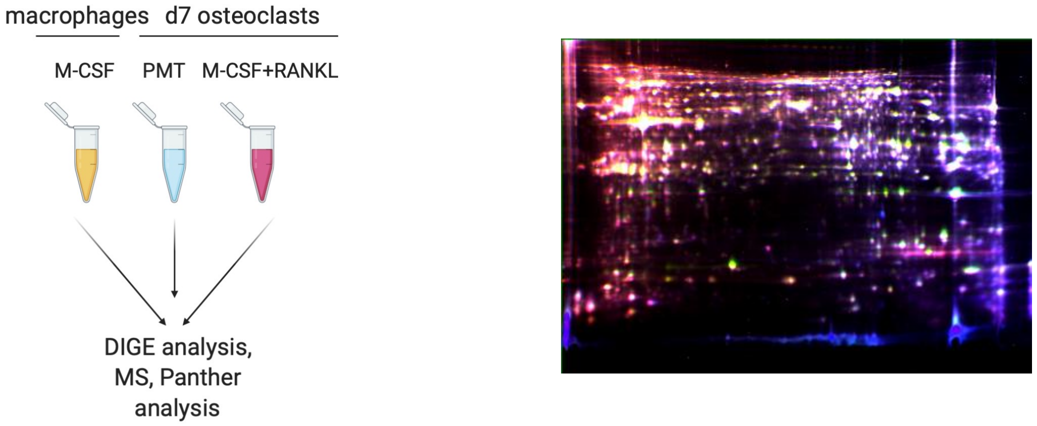


B

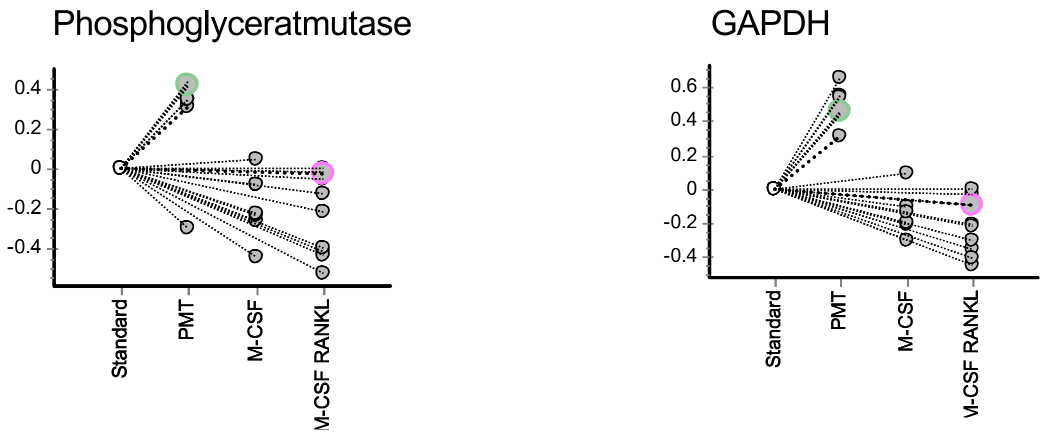


(A) DIGE analysis of the proteome of the different stimulation approaches (M-CSF, M-CSF/RANKL and PMT) of BMDM. Shown is a representative DIGE gel with the overlay of Cy2, Cy3, and Cy5 fluorescent dyes. For the horizontal dimension, a pH gradient from 3-10 was used. In the vertical dimension, the proteins were separated according to their molecular weight (10 kDa-150 kDa). M-CSF/RANKL-treated sample were stained with Cy3, PMT-treated samples with Cy5.

(B) Examples of significant differentially expressed proteins in M-CSF, M-CSF/RANKL and PMT-treated macrophages. Graphic representation of the fluorescence intensity of two significantly altered protein spots, which were evaluated using the BVA module of the DeCyder software. Protein expression was first normalized against the internal standard and then compared between the different samples and gels. Each circle represents the protein spot in the different gels.

Supp Figure 1

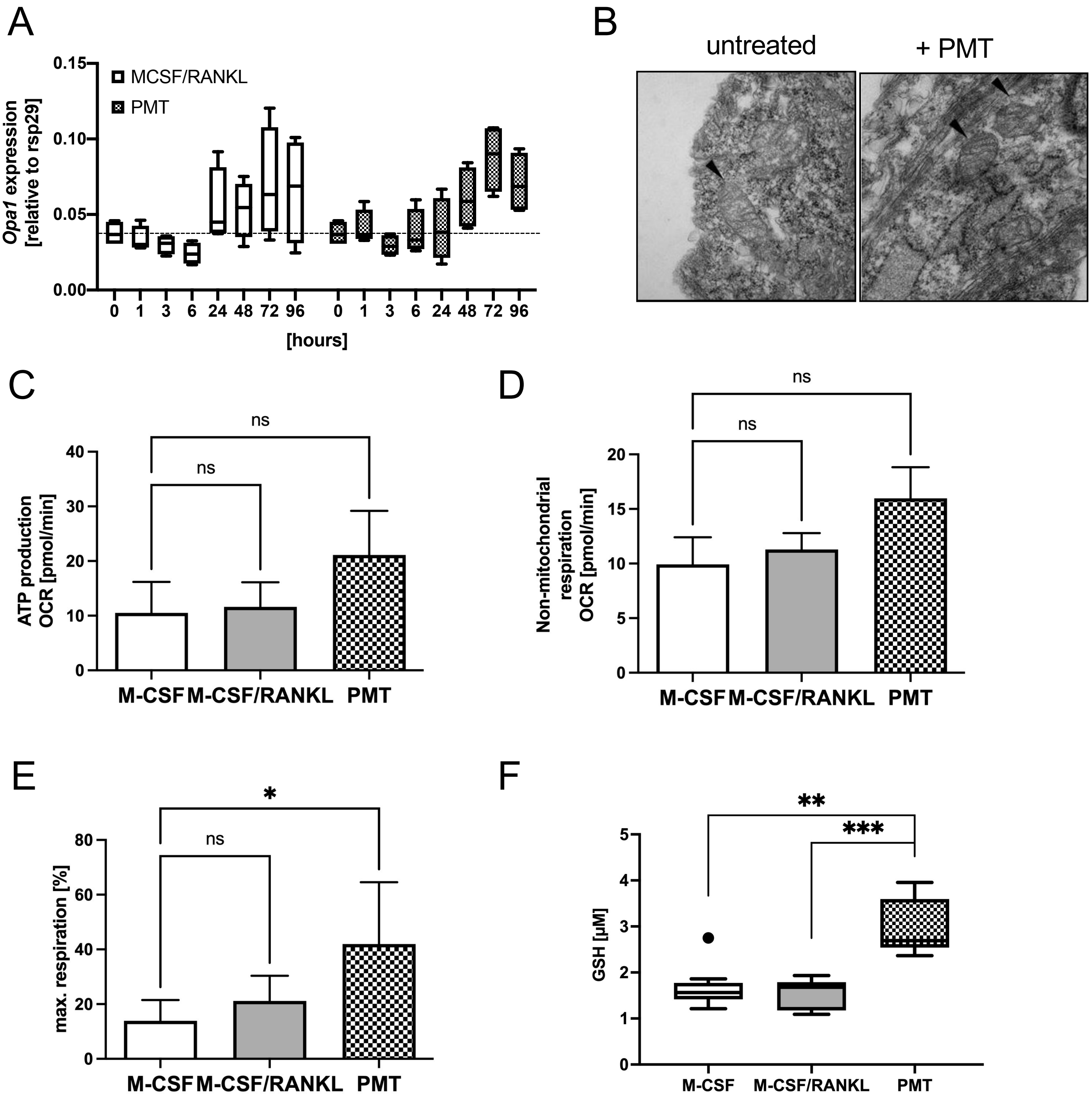


(A) RT-PCR analysis of *Opa1* gene induction in BMDM treated with M-CSF/RANKL and PMT treated cells during osteoclast differentiation. (B) PMT preserves mitochondrial cristae structure in H/R treated H9c2 cardiomyocytes. Examination of mitochondrial morphology was performed by classical electron microscopy. Staining procedure was done by the EM facility of the University of Heidelberg. H9c2 cells were stimulated with PMT or left untreated before fixation with cacodylate buffer and paraformaldehyde and glutaraldehyde. Images were acquired at a magnification of 25k. (C)-(E) Data obtained from Seahorse analysis of mitochondrial activity on day3 of differentiation (Mito Stress Test) (n=3). Quantification of (C) ATP production, (D) no-mitochondrial respiration and (E) maximal respiration. Statistical analysis was done using a Friedman tests for paired samples. (F) Glutathione levels were quantified on day 3 of differentiation and luminescence was measured. concentration of GSH was calculated using a standard curve with glutathione (16 µM to 0.125 µM). One outlier was removed by Tukey analysis. Statistical analysis was done using a Kruskal-Wallis test on unmatched samples.

Supp Figure 2

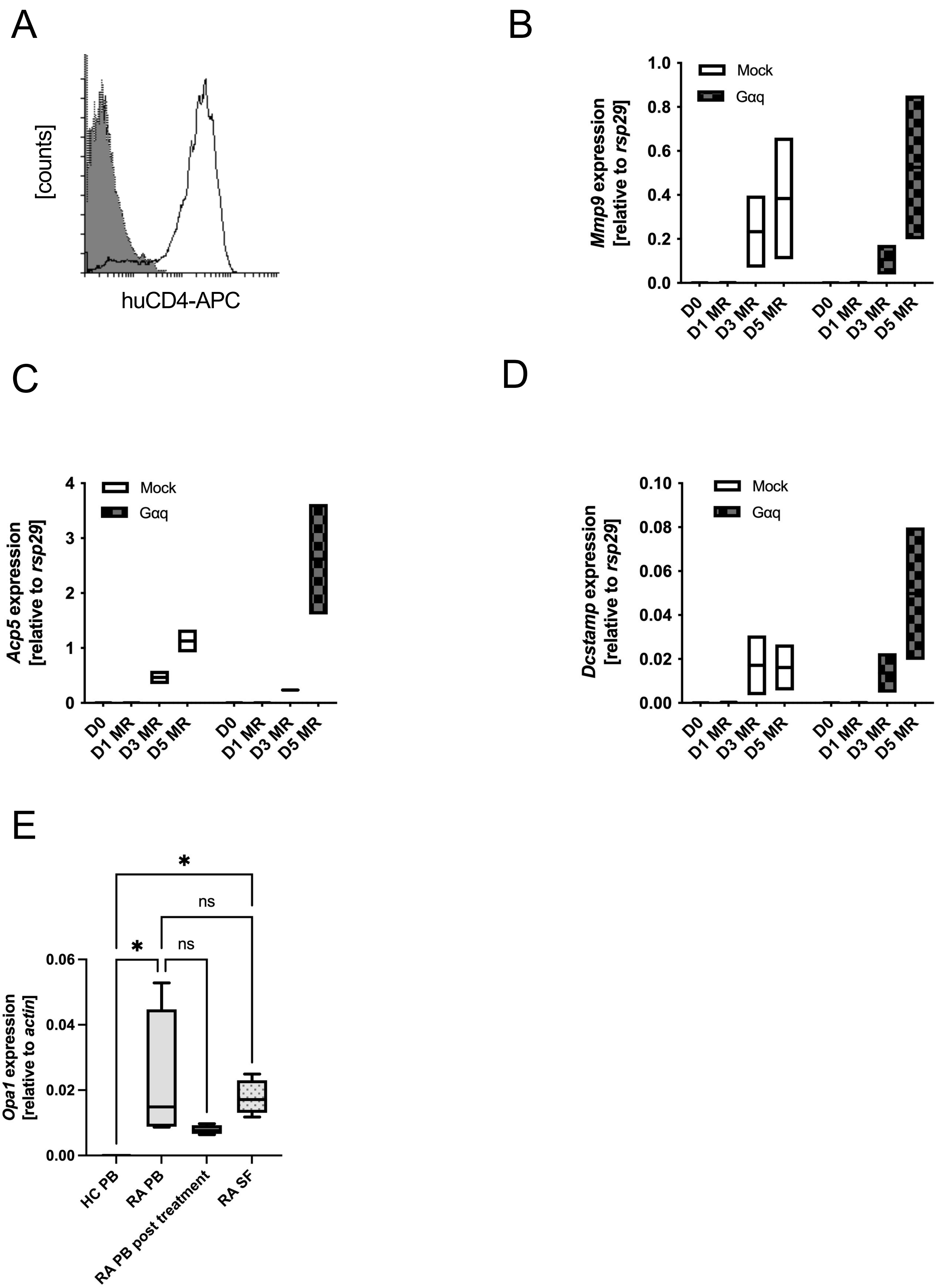


(A) FACS analysis of retrovirally transduced ER-Hoxb8 cells after puromycin selection for huCD4 surface expression. Parental ER-Hoxb8 cells were included as a negative control.

(B) (C)(D) RT-PCR analysis of *Mmp9*, *Dcstamp and Acp5* gene induction in BMDM treated with M-CSF/RANKL and PMT treated cells during osteoclast differentiation (days 0, 1, 3 and5).

(E) RT-PCR analysis of *Opa1* gene induction in human PB samples from healthy donors and RA patients, before and after treatment, as well as SF samples of RA patients.

Supp Figure 3
